# Chemosynthesis enhances carbon fixation and nutrient cycling in an active microbialite ecosystem

**DOI:** 10.1101/2024.12.18.629060

**Authors:** Francesco Ricci, Pok Man Leung, Tess Hutchinson, Thanh Nguyen-Dinh, Alexander H. Frank, Ashleigh v.S. Hood, Vinícius W. Salazar, Vera Eate, Wei Wen Wong, Perran L.M. Cook, Chris Greening, Harry McClelland

## Abstract

Microbialites—carbonate structures formed under the influence of microbial action— are the earliest macroscopic evidence of life. For three billion years, the microbial mat communities responsible for these structures fundamentally shaped Earth’s biogeochemical cycles. In photosynthetic microbial communities, light energy ultimately drives primary production and the ensuing cascade of daisy-chained metabolisms. However, reduced compounds such as trace gases and those released as metabolic byproducts in deeper, anoxic regions of the mat, could also fuel chemosynthetic processes. Here, we investigated the intricate metabolic synergies that sustain microbialite community nutrient webs. We recovered 331 genomes spanning 40 bacterial and archaeal phyla, revealing a staggering diversity fuelled by the biogeochemistry of these ecosystems. While phototrophy is an important metabolism encoded by 17% of the genomes, over half encode enzymes to harness energy from reduced compounds and 12% co-encode carbon fixation pathways, using sulfide and hydrogen as major electron donors. Consistent with these genomic predictions, we experimentally demonstrated that microbialite communities oxidise ferrous iron, ammonia, sulfide and gas substrates aerobically and anaerobically. Furthermore, carbon isotopic assays revealed that diverse chemosynthetic pathways contribute significantly to carbon fixation and ecosystem organic matter production alongside photosynthesis. Chemosynthesis in microbialite communities represents a complex interplay of metabolic synergies and continuous nutrient cycling, which decouples community carbon fixation from the diurnal cycle. As a result, this process mitigates the loss of organic carbon from respiration, enhancing the net productivity of these highly efficient ecosystems.

**Significance:** Microbialite ecosystems are among the most ancient on Earth, having dominated the biosphere for over three billion years and persisting into the present. They serve as critical models for studying past and present Earth-biosphere interactions. In this study, we challenge the paradigm that photosynthesis is the main driver of microbialite primary productivity, emphasizing the fundamental role of chemosynthesis in the global element cycle both in modern extreme environments and throughout Earth’s history. Altogether, our findings provide novel insight into these unique microbial ecosystems, which may have functioned as hotspots for metabolic innovation over geological time.

## Introduction

Complex microbial communities in aquatic environments can form carbonate structures named microbialites. With the earliest fossils dating to around three and a half billion years^1,2^, these structures are representative of the most ancient and persistent microbial ecosystems in Earth’s history. Over Earth’s long history, dense, benthic microbial communities played a pivotal role in shaping the composition of Earth’s atmosphere^3,4^. Notably, they contributed to the rise in atmospheric oxygen following the evolution of oxygenic photosynthesis^3,5^, which amplified global biological productivity by 100-1000 fold^6^. Much of this new productivity first occurred in microbial mats that provided favourable conditions for photosynthetic communities to thrive^7–9^. Additionally, mat communities mediated large fluxes of reduced gases such as carbon monoxide (CO), hydrogen (H_2_), and methane (CH_4_) into the atmosphere^6^. Today, living microbialites are generally confined to extreme environments such as hypersaline lakes^10–14^, where grazing metazoans and plants are typically absent^3^. However, modern stratified microbial communities have a structure resembling those from the Cambrian^15^ and Archaean^5^, positioning these communities as natural laboratories for unravelling the functional processes underlying these ancient systems.

Microbialites are hotspots of microbial diversity^16–18^, yet some key processes that have sustained the ecosystems associated with these structures remain enigmatic. Light is thought to have had a central role throughout their history, initially sustaining anoxygenic phototrophs and later cyanobacteria and algae, driving organic carbon production, carbonate precipitation, and elemental cycling^5,6,19,20^. The resultant photosynthetic end-products shape aerobic niches and fuel complex microbial networks^3^, including by supporting the activity of aerobic heterotrophs^21–23^. Anaerobes dominate beneath the surface of microbial mats, including fermenters, sulfate-reducing bacteria (SRB), and methanogens that catabolise organic matter and gases^6,24,25^. SRB use organic (e.g. acetate, sugars) and inorganic electron donors (e.g. H_2_) while releasing H_2_S. Chemolithoautotrophs can exploit inorganic compounds, encoding genes mediating the oxidation of H_2_, CO, and various sulfur compounds^26–28^, though their ecological and biogeochemical roles within these ecosystems remain poorly understood. While photosynthetic carbon fixation is central to biomass production in these ecosystems^3,29^, this is not the sole process contributing to their primary production. Most notably, the abundant anoxic pockets within microbialites foster ecological niches well-suited for anaerobic autotrophs utilising the Wood–Ljungdahl pathway (WLP)^30–32^. Although gene- and genome-resolved studies have provided insights into the functional potential of microbialite communities, direct activity-based evidence of the complex metabolic interactions that sustain these ecosystems remains limited.

Here we aim to shed light on the primary production and elementary cycling processes sustaining the biodiversity of calcifying microbial mats using living microbialites from West Basin Lake, Victoria, Australia. While it is recognised that microbialite communities have the potential for several photosynthetic and chemosynthetic carbon fixation pathways, we still lack a comprehensive understanding of their mediators, the role of various electron donors in driving these processes, and their relative contribution to primary production. The classic view positions photosynthesis as a leading process driving biological productivity^3,6,33,34^. However, microbialite communities produce substantial amounts of reduced compounds such as H_2_, H_2_S, CO and CH_4_^6,25^, which could fuel diverse chemosynthetic processes. Using living microbialite ecosystems, we integrate genome-resolved metagenomics with high-resolution metabarcoding, biogeochemistry, isotopes, modelling and phylogenetic analysis to quantify the key metabolisms and their contributions in supporting the nutrient web in microbialite communities.

## Results & Discussion

### Taxonomically and metabolically diverse microbes control a complex nutrient web in living microbialites

We deeply sequenced nine metagenomes across three microbialite communities from West Basin Lake in Victoria, Australia. This effort resulted in the recovery of 331 medium-to high-quality metagenome-assembled genomes (MAGs) dereplicated at the species level using a 95% ANI threshold. These MAGs depict a remarkable diversity spanning 40 bacterial and archaeal phyla, including the first genomes from a microbialite community for the phyla JAHJDO01 and UBP6 (Fig. 1). Many of these microbes represent elusive and rare lineages, such as Cloacimonadota, Krumholzibacteriota, Hydrogenedentota, Omnitrophota, and Sumerlaeota. Representatives of these phyla have been previously found in extreme environments including an Antarctic lake^35^, deep-sea trenches^36^ and geothermal springs^37^. The most abundant phyla were Proteobacteria and Bacteroidota, represented by 81 and 56 MAGs respectively (Fig. 1). Cyanobacteria are considered key members of these ecosystems, though we only retrieved one medium-quality MAG of *Halothece* (total relative abundance 3.11%) and 18 MAGs capable of anoxygenic chlorophototrophy (Fig. 1). Millimeter-scale 16S rRNA gene community profiling detected 30 cyanobacterial sequences, which were present at considerably different relative abundance between the analysed samples (sample C: 11.2 ± 7.6 %; sample D: 0.27 ± 0.35 %) (Supp. Data 1). Archaea are also significant members, represented by 18 MAGs spanning Asgardarchaeota, Halobacteriota, Iainarchaeota, Nanoarchaeota, Thermoplasmatota and Thermoproteota (Fig. 1). Eukaryotes have been found in most modern microbial communities^38–40^ and accordingly we identified several major members of diatoms and red and green algae (Supp. Data 1).

**Figure 1.**
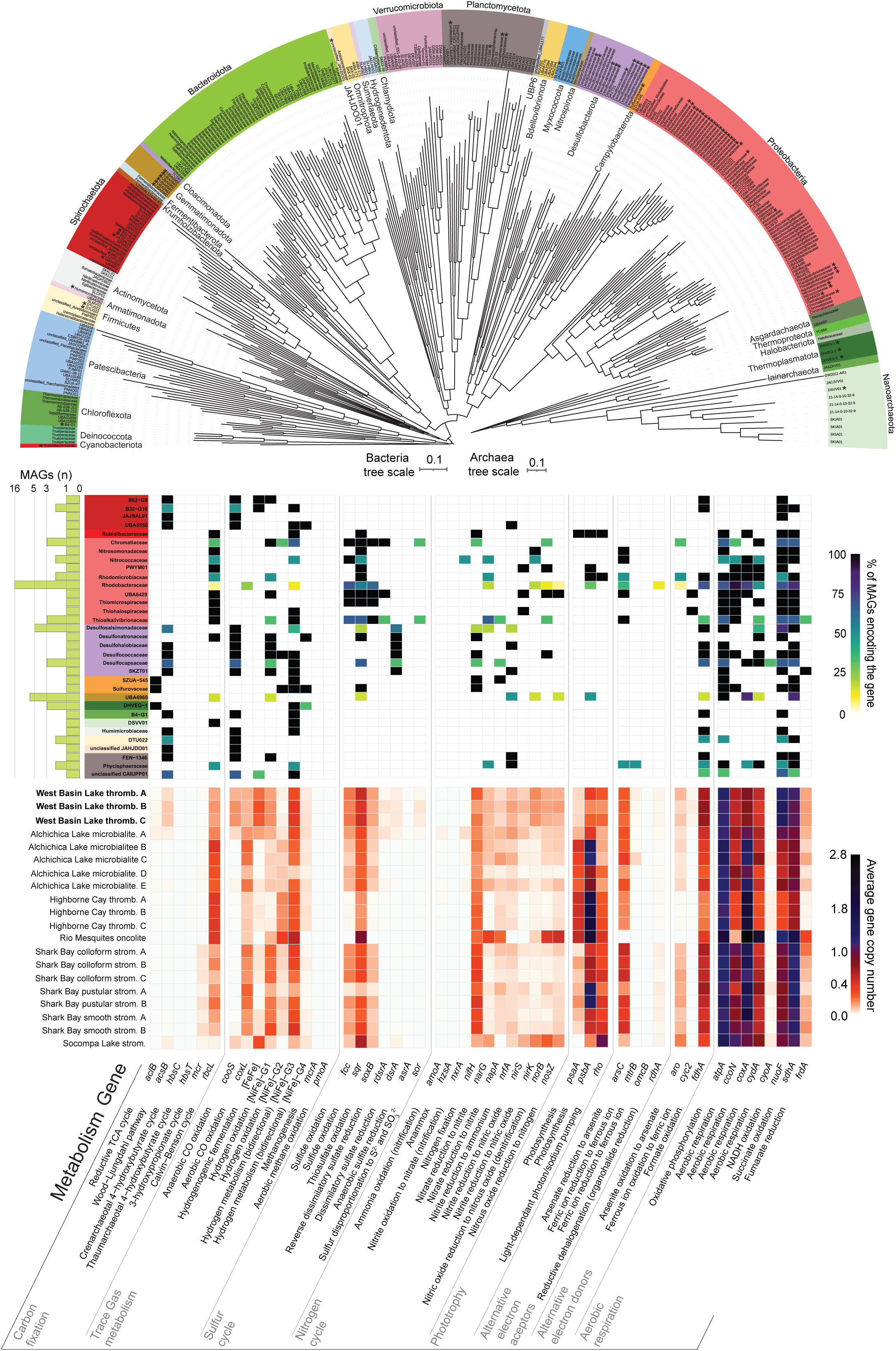
Maximum-likelihood genome trees depicting the taxonomic diversity of 331 archaeal and bacterial metagenome-assembled genomes (MAGs), built with 1000 ultrafast bootstrap replicates using the LG+C10+F+G and WAG+G20 models, respectively. The top heatmap highlights the metabolic potential of microbes co-encoding energy acquisition enzymes with carbon fixation pathways. The side bargraph shows the number of MAGs in each microbial family (GTDB taxonomy). The bottom heatmap shows the abundance of each gene in the metagenomic short reads in West Basin Lake samples (bold) and across 17 publicly available microbialite community metagenomes from five global sites. Homology-based searches were used to identify signature genes encoding enzymes associated with metabolic pathways. To infer abundance, read counts were normalised to gene length and the abundance of single-copy marker genes.

Screening of key metabolic pathways in the MAGs revealed extensive metabolic diversity within West Basin Lake microbialite communities (Fig. 1), potentially underpinning these diverse microbial assemblages. Many microbes appeared to be facultative anaerobes that can input electrons from organic carbon to aerobic and anaerobic respiratory chains with 63% of genomes encoding at least one terminal oxidase (Supp. Data 2). The widespread capacity to oxidise various inorganic compounds (53.2% of the genomes), including hydrogen, sulfide, and carbon monoxide, for supplemental energy likely allows more efficient use of organic carbon for anabolism than catabolism (Supp. Data 2). Furthermore, our data suggest that at West Basin Lake, most organic carbon is produced *in situ*. We identified multiple lines of evidence supporting this hypothesis. First, the lake is not connected to any inlet and lacks aquatic vegetation. Second, we measured moderate concentrations of organic carbon in the water, averaging 4.83 mg/L. Notably, organic carbon abundance increases on average from 3.09 mg/L in shallow to 5.60 mg/L in deeper layers within the microbialites (Supp. Data 3). Third, 13.6% of the generated MAGs encode genes for carbon fixation (Fig. 1). Specifically, 19 MAGs encoded the WLP, whereas 15 MAGs co-encoded the CBB cycle with several other energy harvesting genes (Fig. 1; Supp. Data 2). Some MAGs possess the genetic machinery to utilise multiple energy sources. For instance, a *Thiohalophilus* MAG encodes the CBB cycle with uptake [NiFe]-hydrogenases, sulfide quinone oxidoreductase, thiosulfohydrolase, reverse dissimilatory sulfate reductase and iron oxidising cytochrome (Fig. 1). We also found six MAGs encoding the reductive tricarboxylic acid cycle (rTCA), three of which belonged to the enigmatic group 3 [NiFe]-hydrogenase-encoding Thermoplasmatota class E2 (Fig. 1). Lastly, phototrophy based carbon-fixation via the CBB cycle was mediated by members of the Cyanobacteria, Proteobacteria and Gemmatimonadota (Supp. Data 2) as well as microalgae in phyla Cercozoa, Chlorophyta, Gyrista and Rhodophyta (Supp. Data 1).

To gain a comprehensive, contextual understanding of the metabolic traits supporting energy conservation and primary production in microbialite communities, we conducted a comparative gene-centric analysis of West Basin Lake metagenomes alongside previously sequenced datasets, including thrombolite, oncolite, and stromatolite communities from five global sites (Fig. 1). The capacity for carbon fixation was consistently high across microbialites worldwide. In West Basin Lake, the CBB (*rbcL* av. 19.6%) and the WLP (*acsB* av. 9.92%) were predominant. In the global datasets, the CBB was also the major pathway (32.5 ± 17.0%), while the WLP and 3-hydroxypropionate cycle showed occasional dominance in specific samples (Fig. 1). The potential to utilise inorganic substrates exhibited significant parallels across sites. Sulfide and H_2_ appear to be dominant energy sources, reflected by the widespread oxidation capacity (*sqr*: West Basin Lake av. 60.9%, global 40.1 ± 20.9%; Group 1-3 NiFe hydrogenase: West Basin Lake av. 24.4%, global 22.2 ± 16.0%). The capacity for CO and arsenite oxidation–ancient metabolic traits–was also substantial across all microbial communities (*cooS* & *coxL*: West Basin Lake av. 17.7%, global 11.2 ± 11.7%; *aro*: West Basin Lake av. 11.5%, global 10.1 ± 7.3%). Conversely, photosynthesis genes showed some variation. West Basin Lake had moderate capacity for chlorophylls and bacteriochlorophylls mediated phototrophy (*psaA, psbA:* av. 21.9%) and microbial rhodopsins mediated phototrophy (*rho:* av. 30.5%), whereas the global datasets exhibited broader ranges (*psaA, psbA:* 12.6–141.3%; *rho:* 13.1–113.5%). This variability most likely underscores the influence of local environmental conditions on energy acquisition strategies. Overall, our findings demonstrate that the genetic potential for energy conservation and carbon fixation in West Basin Lake microbialites is broadly representative of other microbial communities forming microbialites that have been studied globally. The high metabolic flexibility observed in these ecosystems enables them to act as efficient engines of biological productivity, supporting diverse and dynamic microbial communities.

### Tight microbial interactions support intricate biogeochemical cycles

Consistent with previous work^13,18,28,41,42^, our findings reveal that microbialite communities have remarkable metabolic diversity, and we sought to resolve the complex molecular interactions underpinning such diversity. Millimeter-scale community analysis revealed high microbial richness throughout the entire microbialite structure (Supp. Fig. 1). While some taxa such as fermenters in the order Clostridiales were confined to deeper anoxic niches (grey layer - Supp. Data 1), many other microbes were distributed across the whole microbialite structure (Supp. Data 1), most notably Cyanobacteriales and the sulfur-oxidiser Campylobacterales (Supp. Data 1). Similarly, members of the obligately anaerobic order Spirochaetales were ubiquitous (Supp. Data 1). These data suggest that the spatiotemporal physicochemical shifts such as the expansion of oxic pockets during the day foster tight metabolic interaction that allow diverse functional guilds to coexist and meet their physiological needs throughout the microbialite structure.

We developed a model to elucidate the array of molecular exchanges occurring within microbialite communities (Fig. 2). Molecular hydrogen (H_2_) emerges as a central molecule. In these communities, H_2_ is primarily produced through fermentation of photosynthetically- and chemosynthetically-derived organic carbon via diverse group 3 [NiFe]-hydrogenases and [FeFe]-hydrogenases, encoded by 90 and 55 MAGs respectively (Fig. 2). The genetic potential for other fermentation pathways in the community is high, with many MAGs encoding marker genes associated with acetate (*acdA, ack, pta*), formate (*fdhA, fdoG, pflD*), and lactate (*ldh*) fermentation (Supp. Data 5). Diazotrophic Cyanobacteria and members of six other phyla (16 MAGs) may also contribute to H_2_ release as an obligate by-product of the nitrogenase reaction. Upon diffusion into aerobic niches, H_2_ is readily utilised by the abundant community of gas oxidisers spanning 25 phyla (106 MAGs; Fig. 2), which may utilise the electrons derived through this process for carbon fixation (Supp. Data 2). Anaerobic processes likely recycle much of the H_2_, including through coupling to denitrification (109 MAGs) and dissimilatory nitrate reduction to ammonium (55 MAGs; Fig. 2). Hydrogenotrophic acetogens were found across each microbialite sample, albeit at low relative abundance (av. 1.2% of mapped reads). H_2_ also fuels methanogenesis and sulfate reduction to varying degrees, leading to the production of methane and sulfide (Fig. 2). Aerobic and anaerobic CO oxidation are significant processes within microbialite communities (34 MAGs; Fig. 2), supporting energy conservation and carbon fixation in eight phyla (Supp. Data 2). A total of 78 MAGs, representing chemolithotrophic and photolithotrophic microorganisms across nine phyla, possess the potential to oxidize sulfide under both aerobic and anaerobic conditions (Fig. 2; Supp. Data 2). This capability likely provides a substantial ecological advantage, facilitating their persistence in environments characterised by dynamic redox fluctuations and intense resource competition.

**Figure 2.**
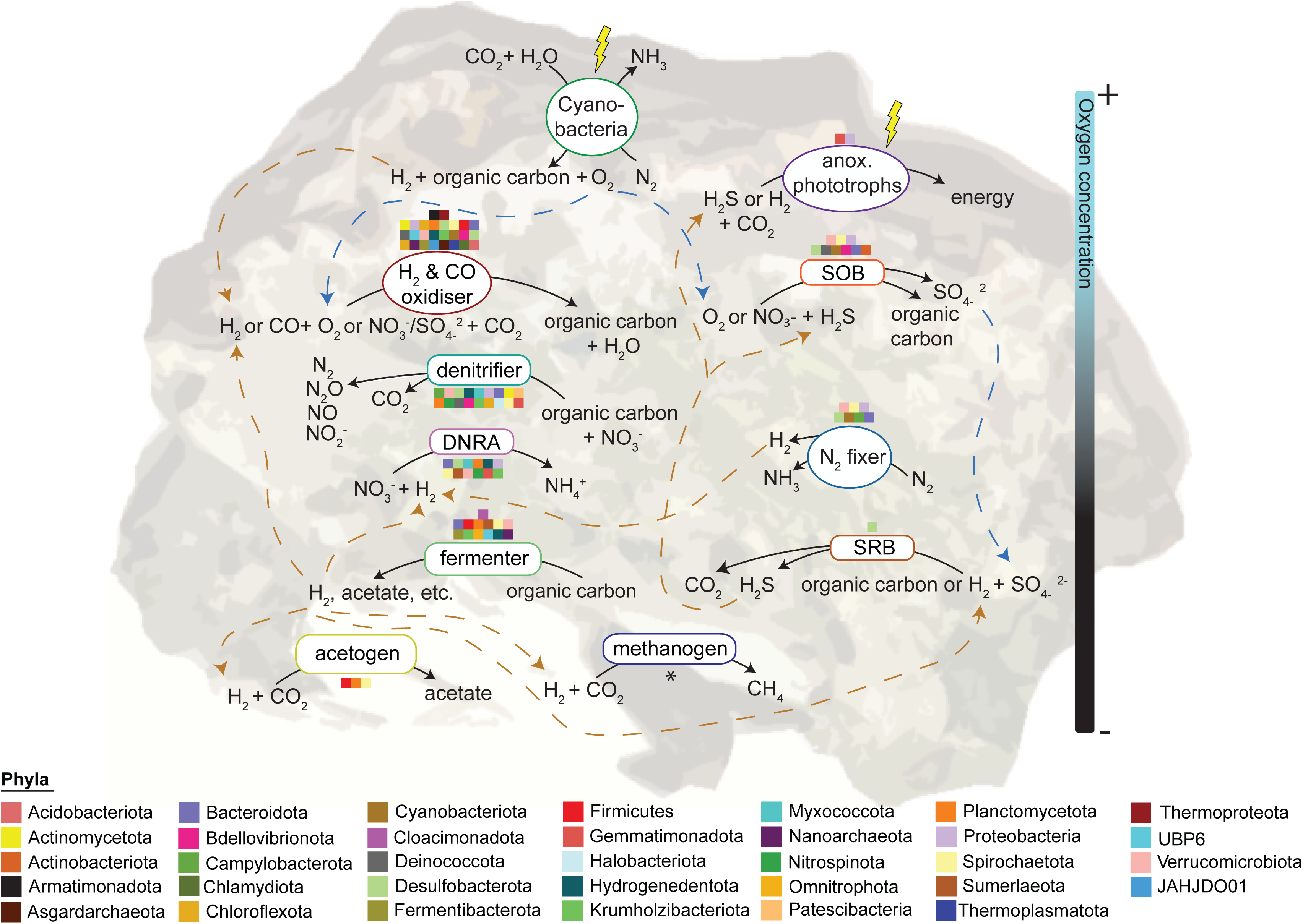
Metabolic model predicted based on genome- and gene-resolved data of the dominant microbial guilds within the microbialite communities of West Basin Lake. This model represents the inferred metabolic pathways and interactions based on the genomic content of these guilds. Blue and brown lines indicate the direction of electron acceptors and donors, respectively. Note that the graphic is an artistic generalization of the data and should not be interpreted as an exact depiction of the microbial community structure. Asterisk (*) denotes that specific metabolic marker genes were exclusively recovered from metagenomic short-read data.

### Microbialite communities mediate broad aerobic and anaerobic elemental cycling

To validate our biogeochemical predictions, we performed *ex situ* microcosm experiments. The potential for oxygenic photosynthesis was assessed using chemical imaging that simultaneously visualises and quantifies O_2_ concentrations in natural samples^43,44^. Algal and cyanobacterial populations, abundant in the microbialite communities, exhibited high activity, producing substantial O_2_ when exposed to light intensities of ∼450 μmol mL^2^ sL^1^, mimicking their natural habitat (Fig. 3; Supp. Fig. 2). The well-illuminated microbialite surfaces demonstrated the highest gross O_2_ production, though this varied markedly both within individual samples (e.g., Sample 2 ranged from 29.31 to 182.26 μmol O_2_ LL^1^ minL^1^) and among samples (Fig. 3; Supp. Fig. 2). Upon transition to darkness, O_2_ was rapidly depleted, primarily through aerobic respiration and diffusion into the overlying water column (Fig. 3). The rapid O_2_ consumption likely contributes to niches for the widespread facultative and obligate anaerobes at the millimetre-scale (Supp. Fig. 2). The O_2_ generated via oxygenic photosynthesis also serves as an electron acceptor for several metabolic pathways, including organotrophy, hydrogenotrophy, carboxydotrophy, and nitrification. Correspondingly, all samples demonstrated oxidation of H_2_, CO, and NH_4_L, albeit at varying rates (Fig. 3).

**Figure 3.**
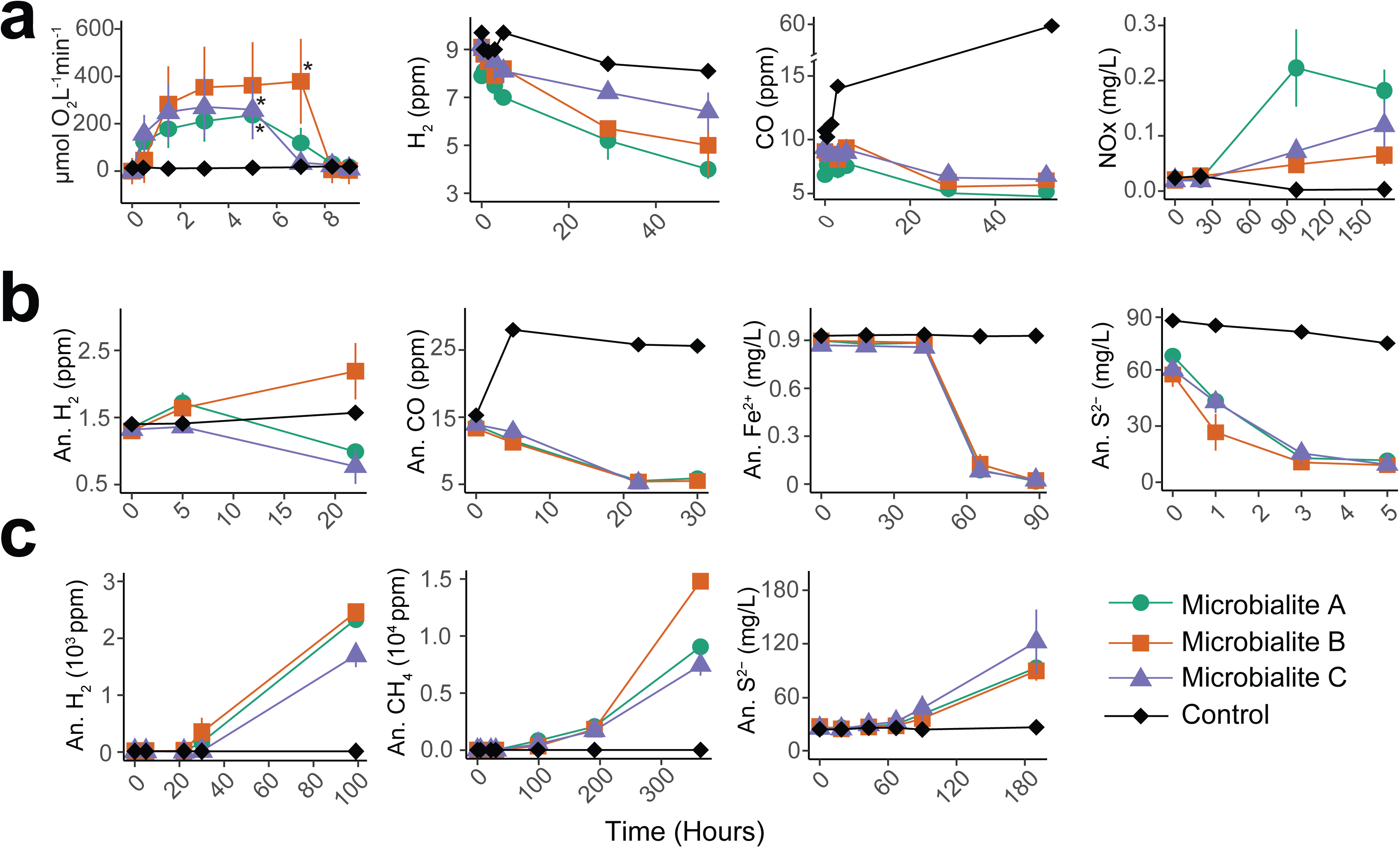
Biogeochemical assays illustrating the metabolic activities of microbialite communities under aerobic (**a**) and anaerobic (**b**, **c**) conditions in microcosms. Oxygen dynamics were assessed using chemical imaging on independent microbialite samples incubated in 4 L glass aquaria, with data presented as the mean ± standard deviation for defined photosynthetic regions of interest. Trace gases, sulfide and ferrous ion measurements were taken in 120 ml sealed serum vial containing 10 g of microbialite slurry and 50 mL of 0.22 μm-filtered lake water. Trace gas microcosms were supplemented with 10 ppm H_2_, CH_4_, and CO in the headspace. S^2-^ and Fe^2+^ microcosms were supplied with either 100 μM NaLS·9HLO (only for consumption) or 6 mM FeClL. All anaerobic microcosms except for S^2-^ production were supplemented with 1.5 mM NO_3_^-^ as an electron acceptor. Nitrification (NOx = NO_2_^-^ + NO_3_^-^) measurements were taken in uncapped 250 mL Schott bottles containing approximately 10 g of microbialite slurry, 100 mL of 0.22 μm-filtered lake water and 100 μM NH_4_ ^+^. In the oxygen plot, asterisks (*) indicate the time when light was switched off, mimicking the onset of darkness. All microcosm experiments were performed in triplicate, with results expressed as the mean ± standard deviation across three replicates. All measurements except for oxygen dynamics were normalised to microbialite slurry wet weight.

We observed consumption of H_2_, CO, Fe^2^L, and S^2^L in nearly all anaerobic microcosms (Fig. 3), confirming the high metabolic flexibility of the anaerobic communities. This metabolic versatility aligns with genomic predictions made by previous studies^13,26,41^ and reflects the capacity of microbialite communities to exploit alternative electron donors to sustain their nutrient web (Supp. Fig. 1). Such flexibility enables them to optimise energy generation across the dynamic physicochemical gradients of the microbialite environment. Consistent with these findings, physicochemical analyses revealed the presence of diverse electron acceptors in the microbialite samples. Specifically, SO_4_ ^2-^ was present in high concentrations (av. 67.85 mg/L), whereas NO_3_ ^-^ at moderate concentrations (av. 2 mg/L; Supp. Data 3).

Our genomic predictions further indicate that microbialites host microbial populations with high capacity for reductive metabolisms (Fig. 1). Microcosm experiments under prolonged anoxia confirmed the production of large amounts of reduced compounds (Fig. 3). Despite methanogens being low in abundance (av. *mcrA* 0.12% of the community), their elevated metabolic rates enable them to contribute notably to CH_4_ production (av. 28.7 ppm h^-1^ g_wet_L¹; Fig. 3), a characteristic previously documented in other environments^45^. Similarly, sulfate reducers were scarce (av. *asrA, dsrA:* 2.44%) but released abundant S^2-^ (av. 7.4 µM h^-1^ g_wet_L¹; Fig. 3). On the other hand, hydrogenogenic fermenters (av. FeFe-hydrogenase: 34.37%; group 3 NiFe hydrogenase: 48.77%) and nitrogen fixer (av. *nifH:* 27.49%) abundance was mirrored in the high H_2_ production (22.7 ppm h^-1^ g_wet_L¹; Fig. 3). These reduced compounds, diffusing through the microbialite structure, fuel a diverse array of aerobic and anaerobic metabolic pathways, thereby supporting microbial energy conservation and carbon fixation within these dynamic ecosystems.

### Chemosynthesis and photosynthesis contribute significantly to microbialites’ carbon fixation

While there’s long-standing evidence for photosynthesis in microbial mats^19,46,47^, recent genomic studies suggest chemosynthetic pathways also facilitate primary production^25–27,48^, though current evidence is fragmentary. To gain a comprehensive understanding of the diversity of microbialite autotrophic populations, we performed phylogenetic analyses of carbon fixation protein sequences. These phylogenetic inferences were further supported by activity measurements using ^14^C incorporation assays conducted with a range of supplemental electron donors, and ^13^C/^12^C organic and inorganic fractionation (Fig. 4d-e). We analysed the phylogenies of protein sequences encoding the acetyl-CoA synthase (AcsB), ribulose 1,5-bisphosphate (RbcL) and ATP-citrate lyase (AclB) (Fig. 4a-c). This analysis revealed an extraordinary diversity of sequences affiliated with at least 16 phyla across the CBB, WLP and rTCA (Fig. 4a-c). These sequences encompassed multiple novel autotrophic clades including two Nanoarchaeota MAGs encoding the CBB, one JAHJDO01 MAG encoding the WLP, and one Polyangia MAG encoding the rTCA (Fig. 4a-c).

**Figure 4.**
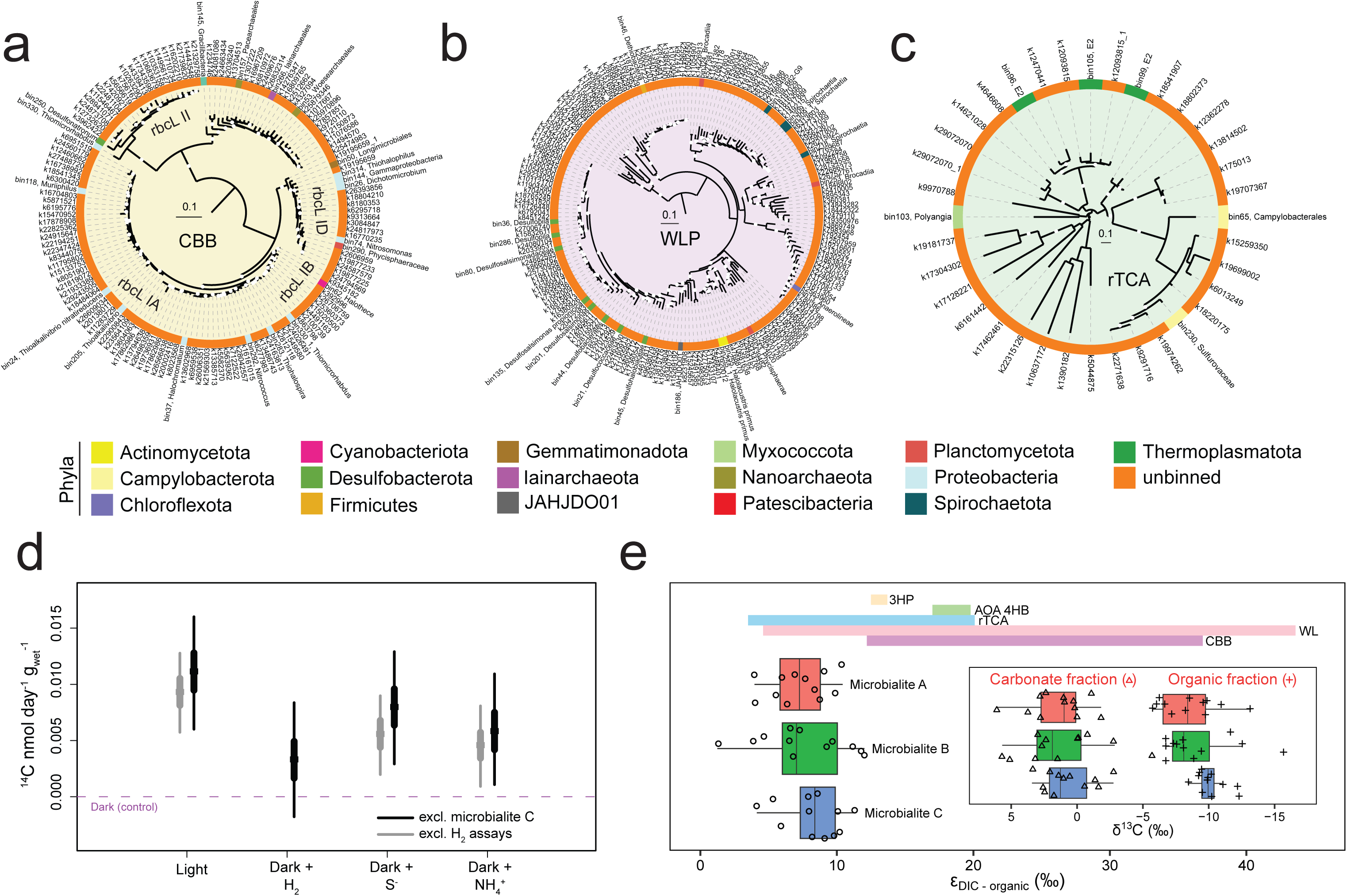
Dominant carbon fixation pathways and activities in microbialite communities. Maximum-likelihood phylogenetic trees of 140 RbcL (**a**), 184 AcsB (**b**), and 36 AclB (**c**) amino acid sequences obtained from the three microbialite samples, constructed using 1000 ultrafast bootstrap replicates. The LG+R5 (**a**), LG+F+I+R6 (**b**), and LG+I+G4 (**c**) substitution models were applied. Sequences derived from binned contigs are highlighted in bold, while those from unbinned contigs are shown without emphasis. Bootstrap support values ≥ 90 are denoted by white circles (**a–c**). (**d**) ^14^CO_2_ incorporation by microbialite microcosms supplemented with different energy sources (light [40 μmol m^−2^ s^−1^], H_2_ [100 ppm], S^2-^ [0.8 mM] and NH_4_^+^ [1 mM]). Credible intervals of effect (electron donors) relative to dark condition denoting 2.5th and 97.5th (thin bar) and 25th and 75th (thick bar) percentiles of posterior probability distribution. Note that the rates do not represent gross carbon fixation rates due to the presence of unlabelled native inorganic carbon (average ∼5.7%; Supp. Data 3) and internally recycled CO_2_ within samples. (**e**) Boxplot showing the carbon isotope fractionation (ε_DIC-organic_ in ‰, Vienna Pee Dee Belemnite standard) between dissolved inorganic carbon (DIC) and organic carbon of three independent microbialites (12 replicates each). Note that the reported fractionation factor of 2.7‰ for aragonite^57^ was used to estimate δ^13^C of DIC from δ^13^C of carbonate fraction, though the primary mineral of sampled microbialites is more likely to be hydromagnesite. Enclosed boxplot shows the natural abundance of carbon isotope compositions (δ^13^C) of the carbonate and organic fractions of the microbialites. Coloured bars depict the range of literature ε_DIC-organic_ values of cellular biomass produced from 3-hydroxypropionate cycle^58,59^ (3HP; yellow), 4-hydroxybutyrate cycle of ammonia oxidising archaea^60–62^ (AOA 4HB; green), reductive tricarboxylic acid cycle^59,63,64^ (rTCA; jade), Wood–Ljungdahl pathway^59^ (WL; pink), and Calvin-Benson-Bassham cycle^53,58,59^ (CBB; violet).

Across the 140 RuBisCO sequences, we recovered four subtypes (Fig. 4a). Notably, RuBisCO subtypes IA-D exhibit intermediate to high specificity for CO_2_ and are generally found across both aerobic and anaerobic habitats, whereas RuBisCO subtype II has lower CO_2_ specificity and is more commonly associated with anaerobic or microaerophilic environments^49^. Consistent with the prevalent anaerobic niches of most microbialites, we recovered 184 acetyl-CoA synthase (AcsB) protein sequences for the WLP (Fig. 4b). However, a large proportion of these sequences most likely were not encoded by autotrophs but rather by members of the phyla Actinomycetota, Desulfobacterota and Chloroflexota that generally utilise the reverse WLP for acetate oxidation coupled with dissimilatory sulfate reduction. Conversely, sequences associated with the Firmicutes, Planctomycetota and Spirochaetota likely represent carbon-fixing acetogens, consistent with prior studies characterizing these phyla^50,51^.

To validate the potential for carbon fixation highlighted through genomic and phylogenetic analysis, we quantified relative chemosynthetic and photosynthetic carbon fixation rates using radiolabeled carbon dioxide (^14^CO_2_). To capture both aerobic and anaerobic carbon fixation processes, we conducted five-day experiments, during which the microcosms transitioned to anaerobic conditions after approximately 24 hours. This transition was inferred based on O_2_ measurements taken simultaneously on other microcosms. ^14^CO_2_ incorporation was detected in all microcosms, except for H_2_-stimulated chemosynthesis in microbialite C (Fig. 4d, Supp. Fig. 3), which most likely resulted from an experimental issue. As anticipated, microcosms exposed to light supported higher ^14^CO_2_ incorporation through photosynthesis (av. 0.0122 nmol dayL¹ g_wet_L^1^) compared to those incubated in the dark (av. 0.00279 nmol dayL¹ g_wet_L¹; Fig. 4d). Dark incubations likely captured the combined activity of anaplerotic processes and baseline chemosynthetic carbon fixation. The addition of electron donors significantly boosted chemosynthetic carbon fixation rates (Fig. 4d). Consistent with the high abundance of genes and MAGs encoding *sqr* and *rdsrA* genes (Fig. 1), S^2-^-supplemented microcosms exhibited the highest chemosynthetic ^14^CO_2_ incorporation rates (av. 0.0084 nmol dayL¹ g_wet_L¹). Similarly, NH_4_ ^+^ and H_2_ supplementation enhanced chemosynthetic carbon fixation (av. 0.00741 and 0.0062 nmol dayL¹ g_wet_L¹, respectively; Fig. 4d).

Finally we probed the origin of West Basin Lake microbialite organic matter through measuring the carbon isotopic composition (δ^13^C) of organic carbon and carbonate fractions. Carbon isotopic compositions of the carbonates (δ^13^C_carb_) were around 1.1 ‰ VPDB (Fig. 4e), consistent with a recent geochemical survey of microbialites^52^. Isotopic compositions of the organic carbon (δ^13^C_org_) were depleted in ^13^C relative to the carbonates, with values ranging from -5.7 to -15.7 ‰ (av. -9.3 ‰; Fig. 4e). However, δ^13^C_org_ is notably more positive than that of organic matter expected from cyanobacterial or algal origin (typically -20 to -30 ‰)^53–55^. It is probable that extensive recycling of organic carbon occurs within the organic-rich microbialite, for example through respiration or fermentation that produces CO_2_ slightly isotopically depleted relative to the carbon of the food source^56^, leading to δ^13^C_org_ enrichment. An alternative explanation is that there is a significant contribution of other autotrophic processes in producing isotopically lighter organic matter. Indeed, in the microbialites of WBL, the degree of fractionation between organic and inorganic carbon (ε_DIC-organic_) is consistent with contributions from diverse C-fixation pathways, most notably rTCA cycle, WLP, and CBB cycles which are associated with larger and smaller isotope fractionations (Fig. 4e). This is in line with the observed high abundance of genes and members mediating these three pathways among the WBL microbialite communities (Fig. 1) and the evidence from ^14^C fixation assay (Fig. 4d). These findings revise the prevailing view that microbial productivity is predominantly driven by photoautotrophs. Instead, we reveal that dissolved inorganic chemical sources and gas substrates are also critical for the primary productivity of these ecosystems, highlighting the significant role of chemosynthesis carbon fixation pathways in supporting stratified microbial communities.

### Conclusions

Through an integrative approach, our study sheds light on the intricate molecular exchanges that underpin the carbon acquisition, energy conservation, and broader nutrient cycling within a model microbialite community, offering a detailed view into how taxonomic and metabolic diversity intersect in such ecosystems. Microbialite communities in the hypersaline West Basin Lake exhibit a metabolic diversity comparable to that observed in other microbialite ecosystems worldwide. This diversity likely arises from the presence of organisms with complementary traits, driven by resource facilitation among community members^65,66^. In microbialite communities, metabolic synergies occur between organisms inhabiting contrasting physicochemical niches^26,67^, which vary on a diel cycle The resultant mosaic of ecological niches align with the physiological requirements of diverse microbial taxa. Consistent with these observations, our analysis simultaneously reveals millimetre-scale overlap of functional guilds throughout the microbialite structure and cycling of key metabolites such as iron, nitrogen and sulfur compounds across their major redox states through intricate molecular handoffs. These processes promote minimal energy loss and enhance ecosystem productivity^68^. In a broader context, the exceptional efficiency of elemental cycling and carbon use efficiency within microbialite communities, combined with the duality of light- and chemically-driven metabolic pathways, suggests that these ecosystems have likely served as hotspots of metabolic innovation throughout Earth’s history.

## Methods

### Characterization of microbial mat organosedimentary structures

The eastern shoreline of West Basin Lake hosts abundant microbialites and living microbial matgrounds^69^. Well-developed microbialite buildups and hardgrounds are exposed on subaerially-exposed benches up to several metres above the current lake level. Previous studies have documented extensive sub-aqueous, well-lithified living microbialites with considerable relief above the lake floor, especially in the adjacent East Basin Lake^69^. The living microbialites documented in this study occur in shallow water of around 0.5 m depth, and are best-developed in benches approximately 1-2 m from the current shoreline in West Basin Lake (Supp. Fig. 4). These living microbial matgrounds appear dark orange-red and have an irregular, undulating surface with a broad-scale relief of several centimetres. In cross section, the mats have a dark orange-red film over a hard to friable, mineralised, cream-white uppermost layer of up to several millimetres in thickness (Supp. Fig. 1). This mineralised layer has an irregular, pustular appearance and is composed of carbonate, likely hydromagnesite^69^. Underneath this, the mat consists of weakly lithified to unlithified mud with several thin colour zones over a several millimetres depth (dark green, purple and grey), followed by up to several centimetres of light green mat (Supp. Fig. 1; Supp. Fig. 4). The sediment texture is commonly clotted and unlaminated. The substrate for the mat is dark, organic-rich mud which may have weak layering preserved through the presence of occasional coarse sediment laminae. Here we use the term ‘microbialite’ to define these microbial matgrounds, as they do form mineralised structures with cm-scale relief above the lake floor. While these modern and recent microbialites are not as extensively developed as the sub-aerially exposed microbialites, they do appear to form significant accretionary structures over time (for example encrusting anthropogenic debris).

### Sample collection, processing and physicochemical parameters

Forming microbialites were collected from West Basin Lake (38° 19’ 24.6468’’ S, 143° 26’ 51.8928’’ E), a hypersaline (salinity 6.5-9.5%) inland crater lake located in Victoria, Australia. Microbialites samples were collected during three field trips in November 2022, March 2024 and May 2024 using pre-sterilized spades and transferred into pre-sterilized 5-gallon buckets containing lake water. At first collection in November 2022, the water temperature was 18.9 °C, dissolved oxygen 90%, pH 8.47 and oxidative reduction potential 2.72 mV. Irradiance at noon was measured using a Walz Universal Light Meter (ULM-500) equipped with a Mini Quantum Sensor (LS-C) at 30 cm and 50 cm below the water surface was ∼650 μmol photons m^−2^ s^−1^ and ∼450 μmol photons m^−2^ s^−1^, respectively. Physicochemical parameters for lakewater and microbialite depth-profile subsamples were measured at the TrACEES Platform, University of Melbourne. The selected physicochemical parameters include total organic carbon, total carbon, inorganic carbon, total nitrogen, sodium, magnesium, potassium, calcium, sulfate and chloride. Nitrate was measured at Water Studies, Monash University.

### 16S rRNA gene sequencing and community analysis

Total DNA was extracted from the layers of two different microbialite samples (three technical replicates per layer) using the DNeasy PowerSoil Kit (Qiagen, Hilden, Germany) on 0.5 g of material. Two sample-free negative controls were also included. Extracted DNA samples were sent to AGRF (Melbourne, Victoria) for library preparation, PCR-amplification and sequencing of the 16S rRNA gene V1-V3 regions on an Illumina MiSeq platform, 2 x 300bp paired-end reads. Sequences were processed using the QIIME2 pipeline v 2022.2^70^. Primer sequences were trimmed using Cutadapt^71^, while DADA2 was employed for merging forward and reverse reads, quality filtering, dereplication, and chimera removal^72^. Taxonomic classification was performed with QIIME2’s feature-classifier plugin. The SILVA v132 QIIME release was used for 16S rRNA gene taxonomy^73^.

### Community DNA extraction and sequencing

Each microbialite sample was homogenised into a slurry. Total DNA was extracted from three different microbialite slurry samples (technical triplicate per sample for a total of nine samples) using the DNeasy PowerMax Soil Kit (Qiagen, Hilden, Germany) on 10 g of materials as per the manufacturer’s protocol. A sample-free negative control was also included. Extracted DNA samples were sent to AGRF (Melbourne, Victoria) for library preparation and sequencing on two lanes using an Illumina NovaSeq SP Flow-cell, 2 x 150 bp for 500 cycles.

### Reads quality control, assembly and binning

Across the three technical replicates of each microbialite sample, we obtained an average of over 16 million read pairs for sample 1, over 56 million read pairs for sample 2, and over 40 million read pairs for sample 3. Reads quality control, assembly and binning were implemented within the Metaphor pipeline^74^. Specifically, raw reads derived from the nine metagenome libraries were quality-controlled by trimming primers and adapters, followed by artefacts and low-quality read filtering using fastp^75^ with parameters *length_required 50*, *cut_mean_quality 30*, and *extra: -- detect_adapter_for_pe*. The nine quality-controlled metagenomes were coassembled using MEGAHIT v1.2.9^76^ with default parameters. Contigs shorter than 1,000 bp were removed. Assembled contigs were binned using Vamb v4.1.3^77^, MetaBAT v2.12.1^78^ and CONCOCT v1.1.0^79^. The three bin sets were then refined using DAS Tool v1.1.6^80^ and de-replicated using dRep v3.4.2^81^ with 95% ANI integrated with CheckM2^82^. Bins completeness and contamination were estimated using CheckM2^82^. After dereplication, we recovered 331 between medium (completeness >50%, contamination <10%) and high-quality (completeness >90%, contamination <5%) metagenome-assembled genomes (MAGs), according to the MIMAG standard^83^. MAG taxonomy was assigned according to Genome Taxonomy Database Release R214^84^ using GTDB-Tk v2.3.2^85^. CoverM v0.6.1 was used to calculate the relative abundance of bins based on the metagenomic reads (https://github.com/wwood/CoverM).

### Functional annotation of binned contigs and unbinned contigs

The sequences of 51 marker genes representing energy conservation, carbon fixation, trace gas metabolism, sulfur cycle, nitrogen cycle, arsenic cycle, iron cycle, formate oxidation, phototrophy, and aerobic respiration were retrieved from the 331 MAGs and unbinned contigs. Open reading frames (ORFs) were predicted using Prodigal v2.6.3^86^, then annotated using DIAMOND blastp^87^ homology-based searches against a custom database^88^ of 51 metabolic marker gene sets described below. DIAMOND mapping was performed with a query coverage threshold of 80% for all databases, and a percentage identity threshold of 80% (for *psaA*) 75% (for *mcr, hbsT*), 70% (for *isoA, psbA, ygfK, aro, atpA*), 60% (*amoA, pmoA, mmoA, coxL, [FeFe], nxrA, rbc, nuoFL*) or 50% (all other databases). MAGs were also annotated using DRAM v1.5^89^.

### Metabolic annotation of metagenomic short reads

Paired-end reads from the nine samples collected in this study and public sequences from Alchichica Lake^28^, Socompa Lake^48^, Highborne Cay^90^, Shark Bay^91^ and Rio Mesquites^41^ recovered from the NCBI SRA under accession numbers PRJNA315555, PRJNA317551, PRJNA197372, PRJNA429237 and MG-RAST 4440067.3 respectively, were stripped of adapter and barcode sequences, then contaminating PhiX and low-quality sequences were removed (minimum quality score 20) using the BBDuk function of BBTools v.36.92 (https://sourceforge.net/projects/bbmap/). Resultant quality-filtered forward reads with lengths of at least 100 bp were searched for the presence of the 51 marker genes described above using the DIAMOND blastx algorithm^92^. Specifically, reads were compared against the custom-made reference databases^88^ of 51 metabolic marker genes for energy conservation, carbon fixation, phototrophy, and hydrogen, carbon monoxide, methane, sulfur, nitrogen, and iron cycling. A query coverage of 80% and an identity threshold of 80% (for *psaA*), 75% (for *hbsT*), 70% (for *atpA, psbA, isoA, ygfK, aro*), 60% (for *amoA, mmoA, coxL, FeFe, nxrA, rbcL, nuoF*) and 50% (for all others) was used. The proportion of community members encoding each gene was calculated by normalizing the gene’s read count (measured in reads per kilobase million [RPKM]) against the average RPKM of 14 universal single-copy ribosomal marker genes.

### Phylogenetic analysis

Maximum-likelihood phylogenetic trees for archaeal and bacterial MAGs were built using GTDBtk^85^ commands *identify* and *align* on high quality MAGs (completeness >90% and contamination <5%). The archaeal and bacterial trees were built using IQ-TREE v2.3.6^93,94^ with 1,000 ultrafast bootstrap^95^ using the LG+C10+F+G and WAG+G20 models, respectively. MUSCLE^96^ was used to align 36 AclB, 184 AcsB and 140 RbcL proteins retrieved between the MAGs and unbinned contigs. AclB, AcsB and RbcL maximum-likelihood phylogenetic trees were built using IQ-TREE v2.2.2.6^93,94^ with 1,000 ultrafast bootstrap^95^ and models LG+F+I+R6 for the AcsB tree, LG+I+G4 for the AclB tree and LG+R5 for the RbcL tree. All trees were plotted using iTOL v6^97^.

### Chemical imaging

The O_2_ sensitive optode preparation included mixing 100 mg of polystyrene, 1.5 mg of indicator (PT (II) meso-tetra(pentafluorophenyl)porphine), 1.5mg of reference (Macrolex yellow®) and dissolved in 1 g of solvent (Tetrahydrofuran) to form a cocktail. The O_2_ cocktails were knife-coated on dust-free polyester foils (goodfellow.com) and the final thickness of the coating was <2 μm. Once dry, the O_2_ sensitive optode was coated with an anti-refractory layer. The anti-refractory cocktail preparation included mixing 100 mg hydrogel D4, 100mg carbon black and 1 g of 100% ethanol. The anti-refractory cocktail were knife-coated on top of the O_2_ sensitive optode and the final thickness of the coating was <3 μm.

The experimental setup included a modified digital single-lens reflex camera (Canon EOS 1000D) with its near-infrared (NIR) blocking filter removed and equipped with a Sigma 50 mm F2.8 EX DG Macro lens. An emission filter (Schott 530 nm, Uqgoptics.com) was fitted to the lens to detect oxygen fluorescence. Following the protocol described by Larsen et al. (RED), O_2_ sensitive optode were excited by four high-power blue LEDs (l-peak = 445 nm, LXHL-LR3C, Luxeon, F = 340 mW at IF = 700 mA) combined with a 470 nm short-pass filter (blue dichroic color filter, Uqgoptics.com). Microbialite sample cross-sections were illuminated using a Schott Leica KL 2500 LCD Cold Light Source. All components were synchronized via a trigger box (https://imaging.fish-n-chips.de) and controlled using the custom software Look@RGB. Each planar optode was calibrated individually in an aquarium maintained at a constant seawater temperature of 20 ± 1°C in a darkened room. The calibration range for the O_2_ sensitive optode was 0 – 360 μmol L^−1^.

All experiments were conducted in a dark room at a constant 20 ± 1 °C to resemble the lake temperature at the time of sampling. Three microbialite samples were cut using a diamond saw exposing their cross-sections. Each of these samples was placed in a 4 L glass aquarium pressing on the O_2_ sensitive optode, which was attached to the aquarium side. Following overnight acclimation in the aquaria, microbialite sample cross-sections were illuminated from above with approximately 450 μmol m^−2^ s^−1^ of light. Irradiance levels in the experimental setup for defined lamp settings were measured using a Walz Universal Light Meter (ULM-500) equipped with a Mini Quantum Sensor (LS-C). Image sequences capturing O_2_ dynamics across the microbialite cross-sections were taken every 5 minutes. Lake water in the aquarium was aerated with an aeration stone connected to an air pump.

Downstream data analysis was conducted using ImageJ v1.53K. Each image was separated into Red, Green, Green2, and Blue RAW TIFF channels. The ImageJ plugin Ratio Plus was used to calculate the ratio of the Red to Green channels (R/G). The resulting ratio images were color-coded using the ‘Fire’ lookup table to visualise O_2_ dynamics. Calibration was performed using the Curve Fitting function, applying an exponential fit with offset for O_2_, based on planar optode calibration values. Brightness and contrast settings were adjusted to display minimum and maximum values of 0 – 360 μmol L^−1^ for O_2_ images. To minimize the influence of water infiltrating in between the O_2_ optode and microbialite cross-section and to quantify microbial O_2_ production, the first image of each experiment was subtracted from subsequent images using the Image Calculator function. To identify the photosynthetic regions on the microbialite sample cross-sections, we overlaid images with the highest O_2_ production onto microbialite cross-section images. The portions of each microbialite sample that showed O_2_ production were identified as regions on interes (ROI). Subsequently values were extracted from the ROI. Control sample in Fig. 3 was an image sequence of ROI measuring a deeper, anaerobic microbialite cross-section portion simultaneously to O_2_ measurements. Proxies for the net photosynthesis (PN) and apparent dark respiration (RD) were calculated by subtracting images taken with a 5 min interval when O_2_ production and respiration were the highest, respectively. Gross photosynthesis was estimated as PG = PN+|RD|.

### Ex-situ biogeochemical measurements

We conducted microcosm experiments to evaluate the aerobic and anaerobic metabolism of microbialite communities. Each experiment was performed in triplicate, with three independent microbialite samples per replicate. Heat-killed samples were prepared using gamma radiation followed by one autoclave cycle at 121 °C for 30 minutes and served as controls. These controls confirmed that the observed element dynamics were attributable to biotic processes.

Aerobic and anaerobic microcosms, including trace gas and S^2-^ and Fe^2+^ additions, were set up in 120 mL serum vials containing 50 mL of 0.22 μm-filtered lake water and approximately 10 g of microbialite to create slurries. The vials were sealed with butyl rubber septa. For aerobic microcosms, the headspace was left with ambient air, while anaerobic microcosms were flushed with helium for 10 minutes to remove O_2_ and then supplemented with 1.5 mM NO_3_ ^-^ as an electron acceptor (except for S^2-^ consumption). The weights of microbialite used in each microcosm were recorded and used to normalise downstream calculations.

Trace gas microcosms were supplemented with 10 ppm H_2_, CH_4_, and CO in the headspace. Sampling of the headspace began immediately after the addition of electron donors and acceptors, with 2 mL of gas extracted at variable time intervals as shown in Fig. 3. In anaerobic vials, the sampled gas volume was replaced with He. Gas concentrations were analysed by gas chromatography using a pulsed discharge helium ionization detector (model TGA-6791-W-4U-2, Valco Instruments Company Inc.), with calibration based on certified standard mixtures of H_2_, CH_4_, and CO (0, 10, 100 ppm in N_2_, BOC Australia).

In anaerobic S^2-^ and Fe^2+^ microcosms, either 100 μM NaLS·9HLO (only for consumption) or 6 mM FeClL was added to helium-purged slurries. At each timepoint, 3 mL of water was sampled and filtered through 0.45 μm pore-size filters. For S^2-^ analysis, 2 mL of the filtered sample was preserved with ZnAc, while for Fe^2+^ analysis, 1 mL was preserved with ferrozine. Both S^2-^ and Fe^2+^ concentrations were measured using a GBC UV-Visible 918 spectrophotometer, following methods described previously^98^.

Microcosms for nitrification were prepared in uncapped 250 mL Schott bottles containing 100 mL of 0.22 μm-filtered lake water, 100 μM NH_4_ ^+^, and approximately 10 g of microbialite sample. At each sampling timepoint, 10 mL of water was filtered through 0.45 μm pore-size filters and stored frozen until further analysis. The filtered samples were analysed for NOx (NO2^-^ + NO3^-^) concentrations using a Lachat QuikChem 8000 Flow Injection Analyzer (FIA) in accordance with APHA methods^99^. For oxygen measurements, microcosms were prepared in 120 mL vials containing 100 mL of 0.22 μm-filtered lake water and approximately 10 g of microbialite slurry. Dissolved O_2_ concentrations were monitored using a FireSting oxygen probe (PyroScience) until the microcosms approached anoxic conditions.

### 14C incorporation analysis

0.25 g of homogenised microbialite sample with 1 mL of filtered lake water (0.22 μm) were prepared in 7 mL scintillation vials with ambient air headspaces. Radiolabeled sodium bicarbonate solution (NaH_14_CO_3_, Perkin Elmer, 53.1 mCi nmol^-1^) was added to a concentration ∼0.1 µM. Triplicates of each sample were prepared and subjected to five different conditions, namely light (40 μmol m^−2^ s^−1^), dark, dark + H_2_ (100 ppm), dark + S^2-^ (800 µM) and dark + NH_4_^+^ (1 mM) and incubated for 5 days. These experiments aimed at capturing both aerobic and anaerobic carbon incorporation metabolisms and according to parallel O_2_ measurements, microcosms likely transitioned to anaerobic after approximately 24 h. After the incubation period, concentrated HCl was added dropwise to each vial and left for 24h with intermittent shaking to ensure excess unbound dissolved inorganic carbon (DIC) was acidified and released as ^14^CO_2_. HCl was added equally to all vials until bubble production ceased and then they were placed at 60 °C under a heat lamp to dry. When dry, 7 mL of scintillation liquid (EcoLume™, MP Biomedical) was added and ^14^C measured on an automated liquid scintillation counter (Tri-Carb 2810 TR, Perkin Elmer). Photosynthetic ^14^C incorporation values were adjusted to account for the photosynthetic ROI areas present on the 0.25g portion of microbialite used in the light treatment. Assuming that the photosynthetic ROI areas identified through chemical imaging represent the microbialite sample regions performing oxygenic photosynthesis, we normalised by applying the ratio of photosynthetic ROI area of microbialite sample 3 which had the largest photosynthetic surface area (photosynthetic ROI sample 1: 3.11%, sample 2: 2.73%, and sample 3: 6.35%) to total microbialite area. We developed a hierarchical Bayesian model to analyze ^14^C incorporation rates under different conditions. The model is implemented in Stan and R (see SI for code). The model consists of N total observations, with I samples, J conditions, and K replicates for each sample condition combination. As the Dark+H_2_ condition experiment failed for microbialite C, we ran two models: one excluding microbialite C and included all conditions (where I=2, J=5, K=3 and N=30), and one excluding the Dark+H_2_ condition and included all samples (where I=3, J=4, K=3 and N=36), so that in each case the model can be fitted to a complete dataset. The likelihood function is given by:

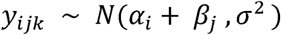

where *α_i_* describes the sample specific effect for the i^th^ sample, β*_j_* describes the condition specific effect for the j^th^ sample, and σ^2^ is the unexplained variance in the data. The Dark condition (j=1) was prescribed to be the control by setting β_1_ _ 0. Weakly informative priors were used. Priors for α*_i_* (*i* ε { 1:*I*}) and β*_j_* (*j* ε { 2:*J*}) were specified as normal distributions with variances an order of magnitude larger than the variance of the total dataset. The prior for σ was a Cauchy distribution. Posterior probability distributions for values of β are presented in Fig. 4d, with Credible intervals defined between the 2.5^th^ and 97.5^th^, and the 25^th^ and 75^th^ percentiles.

### Natural carbon stable isotope measurement

Natural stable isotope measurements were performed to probe possible pathways contributing to the formation of organic matter in microbialites. Three independent microbialites were collected from the same site. For each sample, 12 replicates were prepared by chiselling 12 different regions of the microbialite. Approximately 1.5 g of the chiseled material were dried at 70°C for 2 days, powder-homogenized with a clean mortar and pestle, and further dried at 70°C for 2 days, before shipping to the Bayreuth Center of Stable Isotope Research in Ecology and Biogeochemistry (BayCenSI), University of Bayreuth, Germany for isotopic analyses.

Stable isotope ratios of carbon in the sample are expressed as ratio of the sample compared to the corresponding international standard ratio according to the δ-notation:

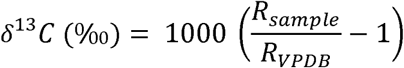

where *R_sample_*is the isotope ratio of ^13^C to ^12^C in the sample and *R_VPDB_*is the ratio in the Vienna Pee Dee Belemnite standard^100,101^.

The relative isotopic composition of the total carbon content in the microbialites was measured using Elemental Analysis-Isotope Ratio Mass Spectrometry (EA-IRMS). To isolate the organic fraction, samples were acidified with one droplet of 85% orthophosphoric acid (Analytical Reagent Grade, Fisher Scientific GmBH, Schwerte, Germany) in silver capsules (5 x 9 mm, IVA-Analysentechnik GmbH & Co. KG, Meerbusch, Germany). The reaction was allowed to proceed at room temperature for at least 48 hours. After this period, the capsules were carefully folded and packed into tin capsules (5 x 12 mm, IVA-Analysentechnik GmbH & Co. KG, Meerbusch, Germany).

The samples were introduced into the oxidation oven of the EA using a helium-purged autosampler. OL gas was simultaneously injected to the carrier gas (helium at a flow rate of 100 ml min⁻¹) to facilitate oxidation in a Fisons-EA-1108 CHNS-O Element Analyzer equipped with a dual reactor setup at 1020°C for oxidation and 650°C for reduction. The GC column (Porapak Q, 80/100, 1.8 m, 2 mm ID) was kept isothermally at 90°C, allowing to isolate the chromatographic peak of COL from accompanying combustion products. The EA was interfaced through an open split (ConFlo IV universal interface, Thermo Scientific, Bremen, Germany) to an IRMS (Delta V Advantage, Thermo Fisher Scientific). The isotopic ratio of the CO_2_ peak was determined through integration and internal calibration against a known reference gas using Isodat 3.0 (Thermo Scientific, Bremen, Germany). After export of the chromatographic areas and isotope ratios; instrument drift, linearity, δ-scale, and the carbon content of the samples was monitored and corrected using a set of external international standards (USGS-62, USGS-63, IAEA-CH7, IAEA-610) spanning the range of measurements.

The fractionation factor (ε_DIC-organic_) between the dissolved inorganic carbon (DIC) and organic carbon of the microbialite samples was calculated using the formula:

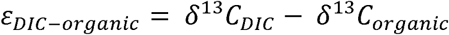

where δ^13^*CDIC* was estimated using a factor of 2.7‰ reported for the carbon fractionation between aragonite and DIC^57^:

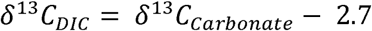

### Statistics and visualization

Downstream statistical analyses were performed in RStudio (version 1.2.5033) using R packages decontam^102^, ggplot2^103^, phyloseq^104^, and vegan^105^. Illustrator v24.0.2 was used for figure editing.

## Data availability

All data supporting the findings of the present study are available. All sequences generated from this work were deposited to the NCBI Sequence Read Archive. BioProject accession numbers for metagenomes, 16S rRNA gene amplicons, and metagenome-assembled genomes are PRJNA1194634, PRJNA1194668, and PRJNA1196970, respectively.

## Conflict of interest

The authors declare no conflict of interest.

## Acknowledgments

We acknowledge Judy and Leon Sjolund for kindly allowing us to conduct our study on their property at West Basin Lake. F.R. was supported by the Early Career Postdoctoral Fellowship (ECPF24-4273843556) awarded by the Faculty of Medicine, Nursing and Health Science at Monash University, and internal funding awarded to H.M. from the University of Melbourne. P.M.L. and A.H. were supported by ARC DECRA Fellowships (DE250101210 to P.M.L.; DE190100988 to A.H.). C.G. was supported by an NHMRC EL2 Fellowship (APP1178715). The isotopic analysis was made available through the BayCenSI Stepping Stones call 2024 awarded to F.R. and P.M.L., financially supported through the DFG-Core Facility Grant with the Project number 461108888 to Prof. Dr. Johanna Pausch. We thank Carina Bauer, Petra Eckert, and Heidi Zier at the Bayreuth Center of Stable Isotope Research in Ecology and Biogeochemistry for their skillful technical assistance. This study used the MASSIVE M3 supercomputing infrastructure.

## Author contribution

F.R., C.G., P.M.L., and H.M. conceptualised the study. Experimental planning and design were conducted by F.R., C.G., T.H., V.E., W.W.W., P.M.L., and P.L.M.C. Fieldwork was conducted by F.R., H.M., and A.H. A.H. provided microbialite description. Data analysis was led by F.R. and P.M.L. Gas chromatography measurements were carried out by F.R., T.H., and T.N., while V.E. and W.W.W. oversaw nitrification measurements. Chemical imaging, oxygen, sulfide and ferrous oxide measurements were performed by F.R. Carbon fixation incubations were performed by T.H. Stable isotope quantification was performed by A.H.F., F.R., P.M.L., and H.M. Metagenome analysis, MAG construction, and annotation were completed by F.R., with extensive bioinformatics support provided by V.W.S. and P.M.L. Resources, supervision, and funding were contributed by C.G., P.L.M.C., and H.M. The manuscript was written by F.R., with input from all authors.

## Supplementary Figures caption

**Supp. Figure 1.**
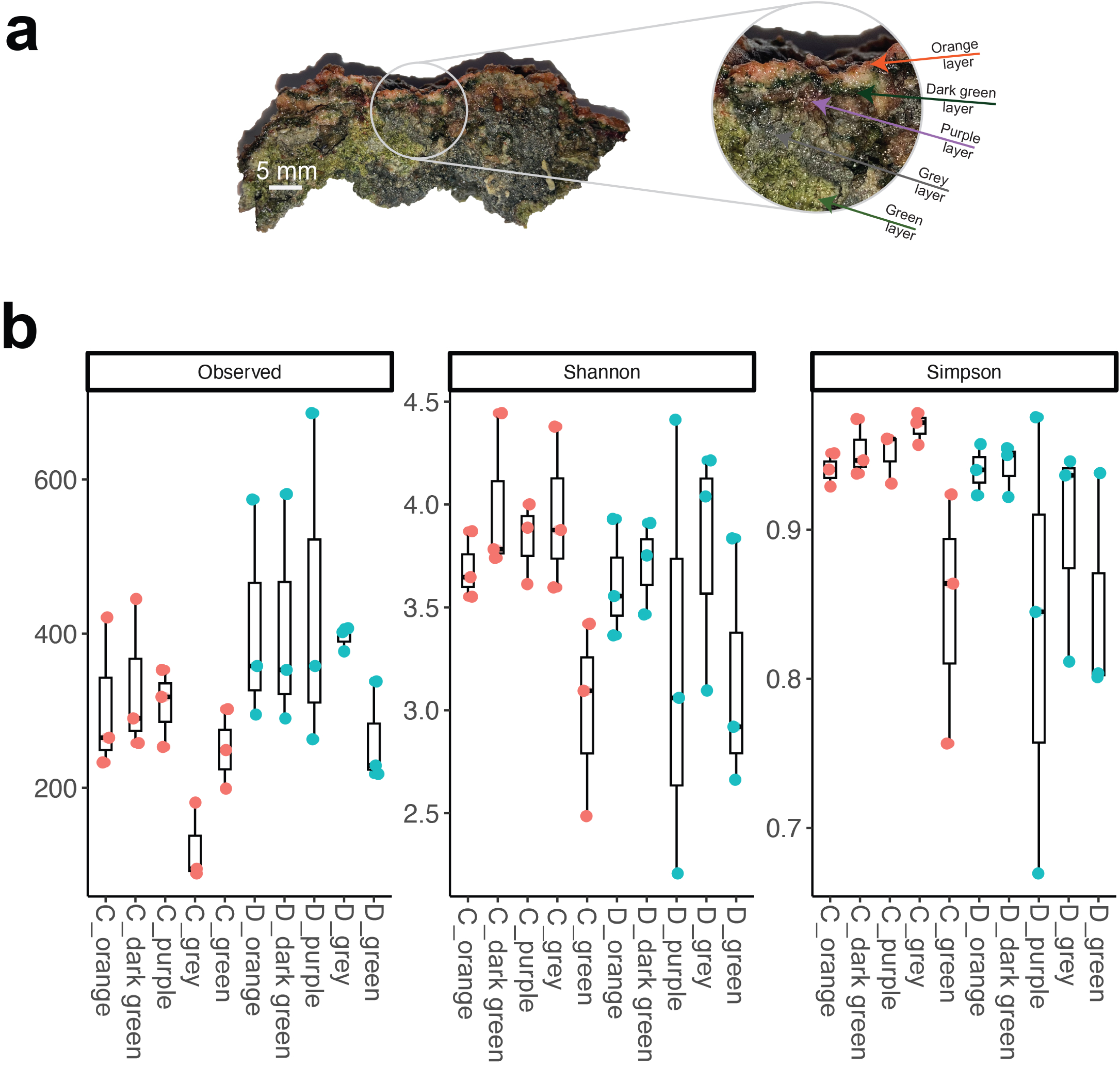
Cross-section of a microbialite with a close-up view highlighting the five sub-samples collected across its structure (**a**). Alpha diversity metrics, including Observed ASVs, Shannon and Simpson, comparing microbial communities from two microbialite samples (**b**).

**Supp. Figure 2.**
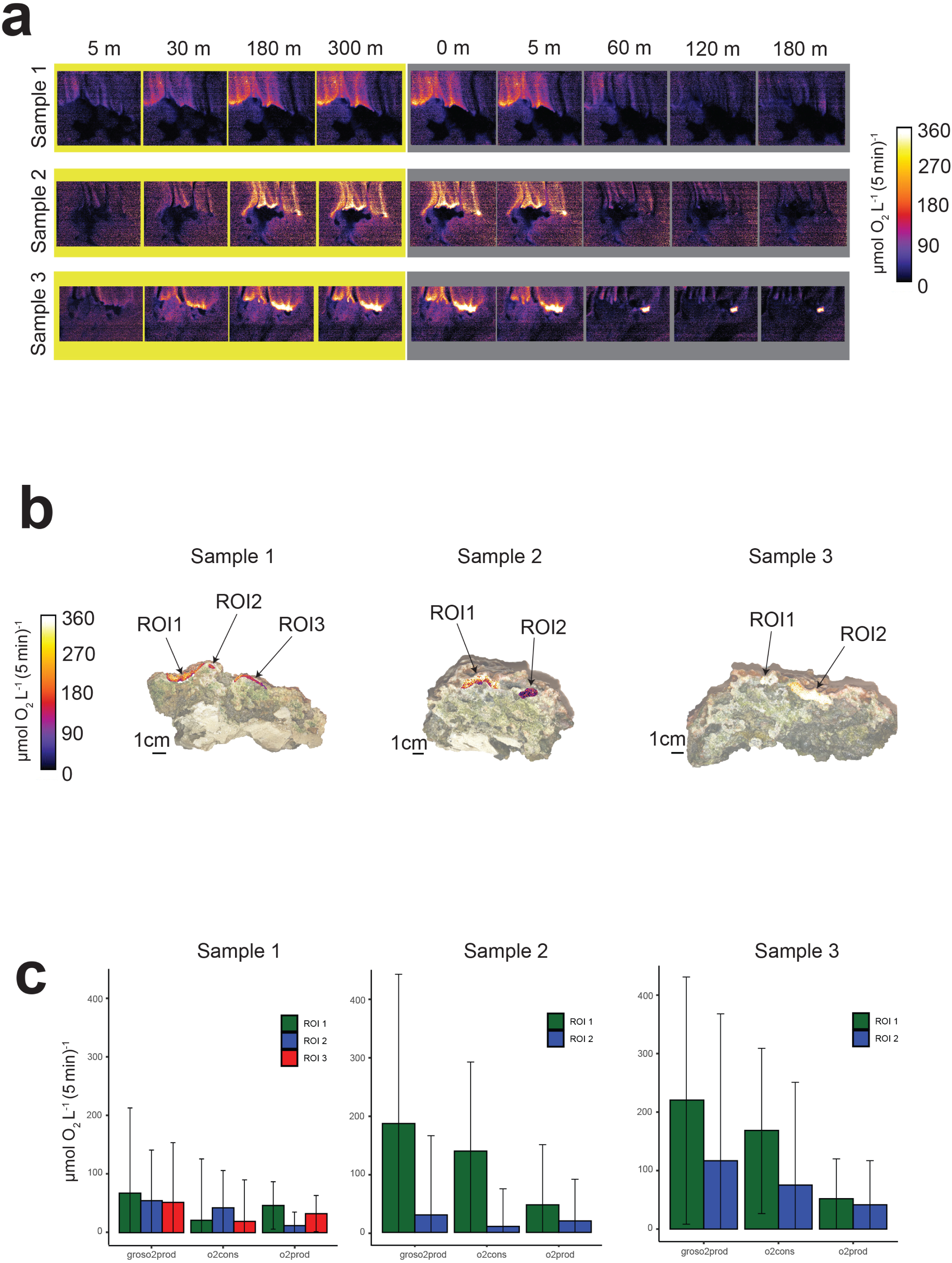
Chemical imaging analysis showing O_2_ dynamics during surficial homogeneous light exposure (yellow background) and after the onset of darkness (grey background) of three microbialite cross-section samples (**a**). Photosynthetic region of interest (ROI) data obtained via chemical imaging, overlaid onto microbialite samples crossed sections (**b**). The ROI colour scale corresponds to the left-hand scale bar indicating oxygen concentration expressed in μmol O_2_ L^-1^ (5 min)^-1^ (**a**-**b**). Bar graphs illustrating rates of gross O_2_ production, O_2_ consumption and net O_2_ production within the photosynthetic ROI of each microbialite sample (**c**). Data are presented as mean ± standard deviation across photosynthetic ROI (**c**).

**Supp. Figure.**
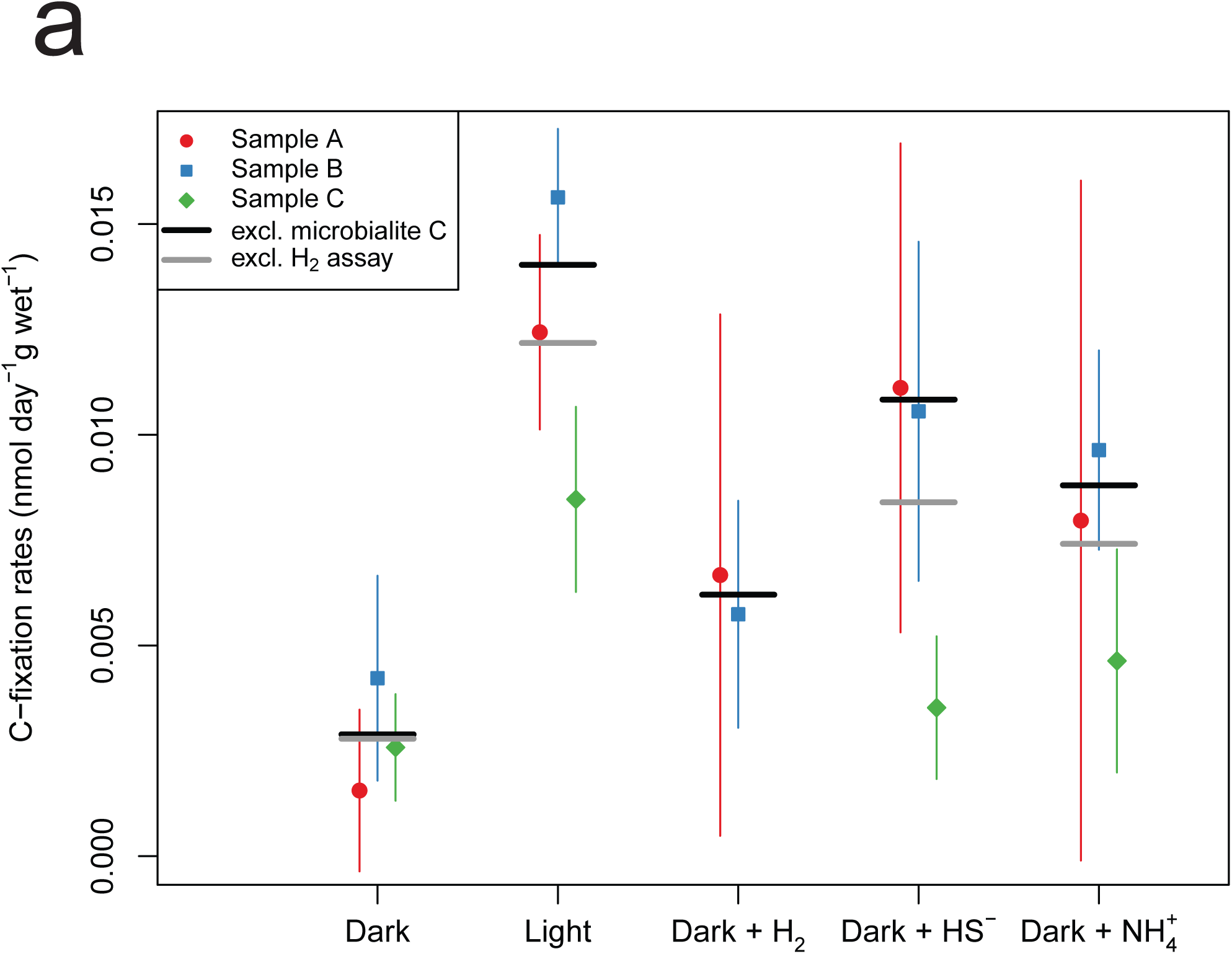
3 Plot illustrating ^14^C incorporation across nine technical replicates of the three microbialite samples exposed to five experimental conditions.

**Supp. Figure 4.**
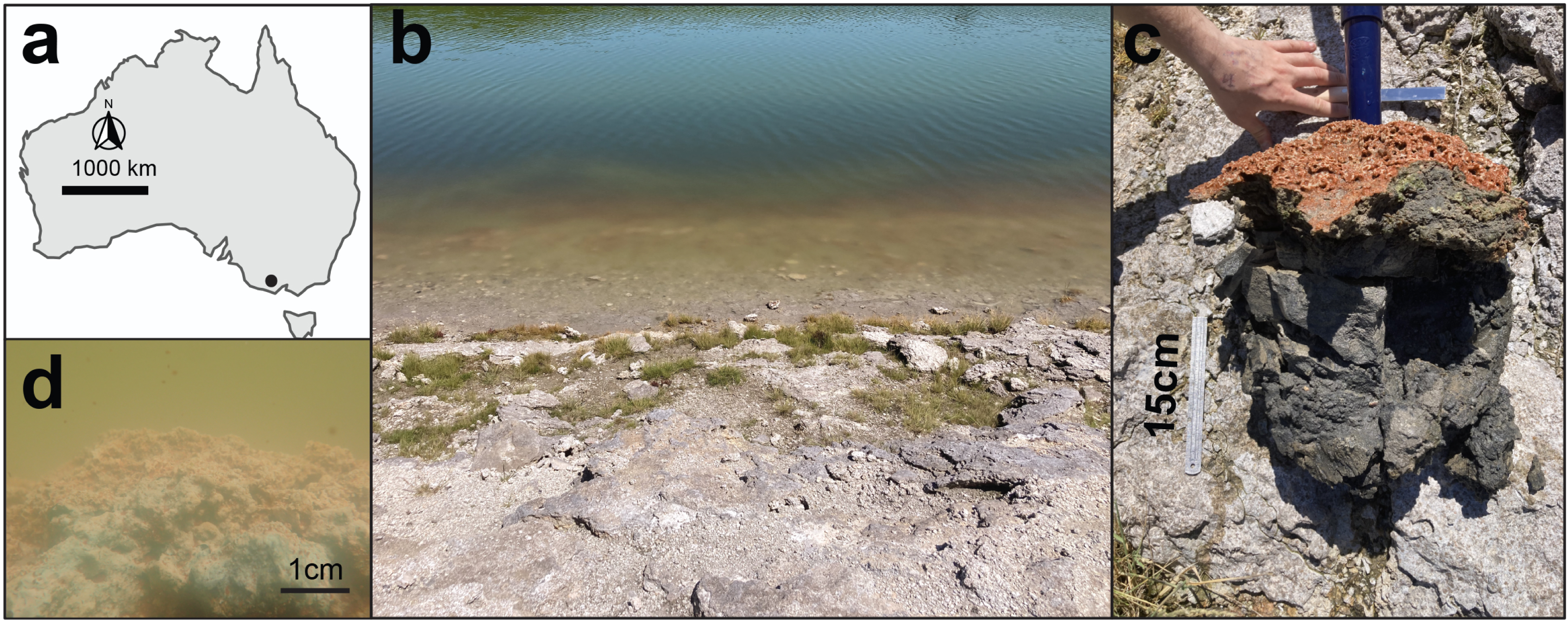
(a) Location of West Basin Lake on Australian map. (b) Picture showing portion of the microbialite reef (in red) at West Basin Lake. (c) example of a freshly collected microbialite. (d) close-up image of a microbialite underwater.

## Supplementary Tables caption

**Supp. Data 1 |** ASV table showing the abundance of bacteria, archaea and eukaryotes in the 16S rRNA gene dataset across the layers of two microbialite samples.

**Supp. Data 2 |** Table highlighting the presence of 57 marker genes across the 331 MAGs generated in this study.

**Supp. Data 3 |** Table showing lake and microbialites physicochemical parameters, namely C, N, Na, Mg, K, SO_4_^2-^, Cl^-^, NO_3_, and ^13^C.

**Supp. Data 4 |** Table presenting metagenomic short read data across three West Basin Lake microbialite samples and 17 publicly available microbialite samples from five global sites.

**Supp. Data 5 |** Table showing genes encoded in pathways involved in acetate, formate and lactate cycling recovered in the metagenome-assembled genomes.

## Notes

https://figshare.com/articles/figure/Microbialite_community_energy_web/28051442

### Competing Interest Statement

The authors have declared no competing interest.

### Summary of Updates

We modified the title to represent better the breadth of the work done in this project, and we added new stable isotope data.

